# Delta activity encodes taste information in the human brain

**DOI:** 10.1101/300194

**Authors:** Raphael Wallroth, Kathrin Ohla

**Author notes:** **Corresponding author:** Kathrin Ohla, PhD, Research Center Juelich, Institute of Neuroscience and Medicine (INM3), Leo-Brandt-Str. 5, 52428 Juelich, Germany.

## Abstract

The categorization of food via sensing nutrients or toxins is crucial to the survival of any organism. On ingestion, rapid responses within the gustatory system are required to identify the oral stimulus to guide immediate behaviour (swallowing or expulsion). The way in which the human brain accomplishes this task has so far remained unclear. Using multivariate analysis of 64-channel scalp EEG recordings obtained from 16 volunteers during tasting salty, sweet, sour, or bitter solutions, we found that activity in the delta-frequency range (1-4 Hz; delta power and phase) has information about taste identity in the human brain, with discriminable response patterns at the single-trial level within 130 ms of tasting. Importantly, the latencies of these response patterns predicted the point in time at which participants indicated detection of a taste by pressing a button. Furthermore, taste pattern discrimination was independent of motor-related activation and other taste features such as intensity and valence. On comparison with our previous findings from a passive (delayed) taste-discrimination task (Crouzet et al., 2015), taste-specific neural representations emerged earlier during this active (speeded) taste-detection task, suggesting a goal-dependent flexibility in gustatory response coding. Together, these findings provide the first evidence of a role of delta activity in taste-information coding in humans. Crucially, these neuronal response patterns can be linked to the speed of simple gustatory perceptual decisions – a vital performance index of nutrient sensing.

## Introduction

The internal representation of sensory events is fundamental to the perception of the external world and adaptive behaviour. Such a representation is achieved in a spatial distribution of neuronal activation which initiates communication across spatially distributed brain areas (cf. Fries, 2015). Rhythmic neuronal activity or oscillations have been regarded as a key mechanism in the process of neural communication in different species (Buzaki, 2006), for instance by sequencing information into temporal processing windows (cf. Lopes da Silva, 1991), and by linking neural assemblies through phase coherence (Tallon-Baudry, 2003; Fries, 2005, 2015). Oscillatory neural activity has been associated with various brain functions (motor action, Salenius and Hari, 2003; consciousness, Ward, 2003; learning and memory, Kahana, 2006; motivation and reward, Knyazev, 2007; attention, Klimesch, 2012) and is well characterized for the sensory systems (visual, auditory, somatosensory, and olfactory) in various species (cf. Koepsell et al., 2010), although the gustatory system is notably absent from this list. Only recently have findings provided evidence of the role of slow-wave synchronized activity in taste processing in rodents (Pavao et al., 2014), whereas no studies at all have characterized the time-frequency dynamics of the human gustatory system.

This lack of knowledge of the frequency by which information is transmitted within the human gustatory system surely does not reflect the importance of the gustatory system. On the contrary, the ability to taste ensures an organism’s survival by enabling the identification of nutrients and avoidance of toxins via a discrimination of taste categories (often referred to as taste qualities). Accordingly, taste categories have been associated with carbohydrates (sweet), electrolytes (salty), acids (sour) or alkaloids (bitter). In rodents, distinct receptors on the tongue respond to chemicals signifying each taste category (Chandrashekar et al., 2006) before the signal is transduced upstream via the gustatory nucleus of the solitary tract in the rostral medulla, and the ventro-posterior medial nucleus of the thalamus to the gustatory cortex in the insula (Carleton et al., 2010). There is evidence of two competing models of taste coding: hardwired, labelled lines with specialized neurons (Chen et al., 2011), and flexible, learning-dependent taste representations (Accolla et al., 2007; Carleton et al., 2010), possibly through broadly tuned neurons (Stapleton et al., 2006). Notwithstanding the unresolved issue of how taste categories are encoded along the peripheral gustatory pathway, at the level of gustatory cortex, however, taste information can be decoded from dynamic activity patterns obtained from neuronal ensembles (Jones et al., 2007) and local field potentials (Pavao et al., 2014) in rodents, and large-scale EEG scalp recordings in humans (Crouzet et al., 2015). The availability of taste information from large-scale recordings enables investigations of the time-frequency dynamics of cortical information transfer in taste perception.

For our investigation, we recorded multi-channel head-surface electroencephalography (EEG) in human participants while they detected salty, sweet, sour, or bitter solutions, in order to investigate the neural mechanism by which the human gustatory system encodes taste information. First, we investigated whether the taste-evoked electrophysiological response would selectively engage a specific frequency band, given that frequency-specific neuronal signatures have been observed in other sensory systems (Koepsell et al., 2010). Since participants received four different tastants, we were further able to test whether taste-specific content is represented in the frequency-specific activity. Second, because such an observation may not be a mere by-product of network activity, but bear functional relevance for perceptual decisions (Harmony, 2013), we hypothesized that the timing of the neural gustatory response would predict the timing at which participants detect a taste. Third, because task dependency has been reported with respect to taste-related behavioural responses in humans (Halpern, 1986; Bujas et al., 1989) and cortical activation in rodents (Fontanini and Katz, 2009), we probed the flexibility of human gustatory processing by comparing taste-evoked neural responses between the speeded detection task presented here and a previously reported delayed categorization task (Crouzet et al., 2015).

## MATERIALS AND METHODS

### Participants

Sixteen healthy participants (12 women; mean age 28±5.1 years; BMI 22±3.0) completed the study. Participants reported having no taste impairments and no history of neurological or psychiatric disease. They signed informed consent prior to the start of the experiment and received monetary compensation for participation. The study protocol conformed to the revised Declaration of Helsinki and was approved by the ethics board of the German Psychological Society.

### Stimuli

Tastants were prepared as aqueous solutions with a clear taste by dissolving 3.8 g salt (salty; NaCl, from a local supermarket), 0.75 g citric acid (sour; SAFC, CAS#77-92-9), 15 g sugar (sweet; sucrose, from a local supermarket), 0.08 g quinine monohydrate (bitter; Honeywell Fluka^®^, CAS#207671-44-1) and 2 g Splenda^®^ (sweet; Tate & Lyle) in 100 ml of distilled water. Splenda trials were not included in the present analysis. Taste stimuli were 210 μl aliquots embedded in a regular stream of water pulses and delivered over a 900 ms period in the form of aerosol pulses to the anterior part of the extended tongue (Crouzet et al., 2015). Importantly, participants kept a slightly opened jaw and an extended tongue during the experiment, which inhibits oromotor behaviour such as tongue, lip or jaw movements and requires no swallowing of the liquids. Notably, this protocol further prevents innate craniofacial responses of the lower facial area, which are known to vary between non-sweet tastes (Rosenstein and Oster, 1988).

*Experimental procedure.* Each trial started with a central black fixation cross on gray background presented for a variable amount of time (0.8-1.5 s) on a 24” thin film transistor (TFT) screen placed at a distance of 45 cm in front of the participant. The fixation cross instructed participants to refrain from any movements; it remained on screen during taste stimulation (0.9 s) until the end of the response-time measurement (2.1 s). Participants were instructed to press a button on a serial response box (Psychological Software Tools, Inc.) with the right or left index finger as soon as they tasted anything (simple response-time task). Response hands were switched between blocks and the starting hand was counterbalanced across participants. The fixation cross was then replaced by the instruction to rate the intensity and pleasantness of the taste, by moving the mouse cursor along a visual analogue scale that was labelled at the end points with “no sensation” and “extremely intense” and “extremely unpleasant” to “extremely pleasant”, respectively. The rating procedure lasted 8 s and was followed by a blank screen for 9 s to minimize adaptation. Overall, 300 taste stimuli (60 per tastant) were presented in pseudo-random order over the course of six experimental blocks. The experiment lasted approximately 120 minutes including breaks between blocks and an initial training during which participants were familiarized with the procedure. Participants were presented with brown noise via noise-isolating in-earphones to mask any auditory cues from the spray pulses.

### EEG data acquisition and pre-processing

Participants were seated in a sound-attenuated recording booth (Studiobox GmbH, Walzbach, Germany) with the gustometer positioned outside to minimize auditory cues from the device. The electroencephalogram (EEG) was recorded with an activCHamp amplifier system (Brain Products GmbH, Munich, Germany) at a sampling rate of 500 Hz with analogue 0.01 Hz highpass and 200 Hz lowpass filters using PyCorder (Brain Vision LLC, Morrisville, NC, USA) and with 64 Ag/AgCl active electrodes placed in an elastic cap according to the extended 10-10 system. The EEG data were processed offline using EEGLAB (Delorme and Makeig, 2004) in MATLAB (Mathworks, Natick, MA, USA) and Autoreject (Jas et al., 2017) in Python. First, we resampled the data to 200 Hz to improve the signal-to-noise ratio and to reduce computation times (Grootswagers et al., 2017). We removed slow drifts with linear detrending and line-noise (50 Hz in Germany) with a set of multi-tapers over sliding time windows implemented in the CleanLine plugin. Second, we segmented the continuous data into epochs from –1.5 s to 2.5 s relative to stimulus onset and applied Autoreject for noisy channel interpolation and epoch exclusion. Less than 1% of all trials were removed. Third, we computed an extended Infomax independent component analysis implemented in EEGLAB (ICA; Makeig et al., 1997) to identify artefactual components with manual inspection and ADJUST (Mognon et al., 2011), which uses temporal and spatial characteristics of the independent components to detect outliers. Components representing common ocular, cardiac or muscular artefacts were subtracted from the data. Lastly, the data were re-referenced to the average of all channels.

### Experimental design and statistical analysis

We employed a within-subjects design with one factor (taste category) with four levels (salty, sour, bitter, sweet). All analyses were conducted within participants and are described in more detail in the respective subsections below. The wavelet transformation was performed in MATLAB using EEGLAB, decoding was implemented with custom scripts in R (R-Core-Team, 2017).

### Time-frequency decomposition

To obtain a time-frequency representation of the EEG data, we computed the spectral power and phase angles via continuous Morlet wavelet transforms of single trials for the frequency range from 1 to 100 Hz in 40 logarithmically spaced steps). We increased the length of the wavelets linearly from 1 cycle at 1 Hz, to 15 cycles at 100 Hz to optimize the trade-off between temporal resolution at lower frequencies and spectral precision at higher frequencies. The convolution was performed non-causally (i.e. a time point is the center of the time window) in steps of 10 ms from –0.2 to 1.5 s. The result of the convolution is a complex number of which the magnitude represents the spectral power and the angle the phase. For visualization, we quantified the degree of event-related phase synchronisation across trials with the inter-trial coherence (ITC; cf. Busch et al., 2009), which takes values between 0 (no synchronisation) and 1 (perfect synchronisation).

### Taste discrimination analysis

To search for taste information in the time-frequency spectrum, we performed a multivariate pattern analysis (MVPA, cf. Kriegeskorte, 2011) at every time and frequency step, using the log-transformed spectral power and phase angles in radians. The MVPA was implemented with a L2-regularized linear support vector machine (SVM; Fan et al., 2008) within subjects in a stratified leave-one-trial-out cross-validation (CV; i.e. on every iteration, a trial of each taste is left out) for optimal model estimation. The multi-taste classification problem was solved via pairwise binary classifiers with the final performance as the average over the six one-versus-one comparisons (cf. Hand and Till, 2001). Thus, the decoder would see the instantaneous topographical distribution of the spectral power or phase angles of a large subset of trials and learn the taste-specific systematicity in their patterns. The availability of such taste-specific information was tested by generalizing these patterns to the taste-evoked responses of the left-out trials.

We optimized the regularization constant C in incremental steps of negative powers of 10 (10^-4:0^) in a nested CV. In this tuning protocol, the training set was iterated in an inner 10-fold stratified CV to search for the C which achieved optimal performance across time points on the inner subset of the data. The classifier was then trained on the full training set with the best value for C of the inner tuning set and tested for its performance on the left-out trials which were neither part of training nor tuning. We evaluated the classifier performance with the area under the receiver operating characteristic curve (AUC), which provides a balanced accuracy metric (50% performance corresponds to chance). For the assessment of statistical significance (*p* < .05) and adjustment for multiple null hypothesis testing, we performed cluster-based permutation analyses on the AUC scores (Maris and Oostenveld, 2007).

### Task dependency of taste coding

After identification of the frequency band that contained taste information, we performed an MVPA which incorporated both power and phase characteristics to compare taste information coding between the task reported in the current study (hereafter, active or speeded taste detection) with a previously reported task, during which participants were to taste the stimuli and to discriminate tastes only after presentation (hereafter, passive or delayed taste discrimination; Crouzet et al., 2015). For this, we isolated the signal via zero-phase Hamming-windowed sinc finite impulse response (FIR) filter (–6 dB cutoff, maximum passband deviation of 0.2%, stopband attenuation of –53 dB). Epochs were resampled to 100 Hz to match the temporal resolution of the wavelet analysis, and filtering was performed on epochs from –1.2 s up to 2.5 s to minimize aliasing before reducing the epochs to the –0.2 to 1.5 s interval of interest. The decoding procedure was performed as described above (see Taste discrimination analysis). Statistical significance of above-chance performance was assessed via one-sided binomial tests and adjusted to a minimum of 100 ms of *p*-values < .05. Differences between the decoding performance scores for the two tasks were compared via two-sided Wilcoxon rank sum tests and adjusted to a minimum of two consecutive time points.

### Controlling for the influence of motor activation

To ensure that the classifiers were not using information related to the motor response in order to discriminate between the tastes, we ran a control analysis in which we repeated the multiclass MVPA with a careful separation of response sides. Here, we trained the classifiers to discriminate between tastes by means of the electrophysiological patterns of trials with a left button press, and tested their performance on patterns of trials with a right button press, and vice versa (out of sample classification). If the classification performance was comparable to the performance found for taste decoding this would demonstrate that the classifier is independent of motor activation and that it indeed used taste-specific information.

### Correspondence between neural and behavioural responses

To test whether the neural taste signal predicts taste detection times, we estimated signal onsets for single trials. We performed the MVPA within a taste category to discriminate a trial against its water baseline (average of 200 ms prestimulus). This analysis is ideal for estimating the classification performance independently of the contrast to the other tastes, and the decoding task closely resembled the taste-detection task that participants performed. We applied the same CV and tuning protocol as for the previous analyses. Trials without responses or with responses faster than 100 ms were excluded (3.5% of trials). To determine the time at which taste patterns emerged at the level of single trials, we used a strict adjustment procedure. To account for cases in which the correct label may be assigned by chance, we estimated for each correct decision a decoder’s confidence through Platt scaling (Platt, 1999) and computed the lower 5^th^ percentile of this confidence estimate for each C parameter (regularization shrinks confidence). The onset within a trial (i.e. the emergence of a pattern which successfully distinguishes a taste from water) was determined as the earliest point before a motor response at which the decoder’s decision was correct for at least 100 ms and with a confidence level above the (C-dependent) 5^th^ percentile threshold. Trials without such a streak of correct classification were removed from the analysis (17.4% of trials). We computed hierarchical linear mixed regression models to examine the link between the estimated onsets as the predictor of response times. We defined the intercepts as random effects at the second level (participants), and the intercepts and slopes as random effects at the third level (tastes). To examine whether the emerging patterns could be linked to other taste features besides taste identity, we also computed analogous models with participants’ rating scores of taste intensity and pleasantness as the dependent variable. We report the fixed effects with their standard errors and estimated *p*-values, using (*χ*^2^-distributed) likelihood ratio-tests against the restricted (intercept-only) null models.

## RESULTS

### Taste information encoded by delta activity

To investigate the encoding of taste information at a specific frequency, we used SVM classifiers at each time and frequency step to decode the four tastes using either the spectral power (Fig. 1A) or the phase angles (Fig. 1C). We found significant clusters of increased classification performance in the lower part of the frequency spectrum. The classifiers decoded taste-specific information from the activation patterns of the spectral power in a major cluster largely contained within the 1 to 4 Hz band (30 to 1160 ms; AUC_max_ = 56.4%±1.2, *p* < .0001, at 1.4 Hz and 360 ms; Fig 1B), which extended into the 8-12 Hz range between 480 and 860 ms. Similarly, decoding of the phase angles revealed a major cluster contained within the frequency spectrum below 8 Hz (10 to 1260 ms; AUC_max_ = 64.3%±2.0, *p* < .0001, at 2.9 Hz and 300 ms; Fig 1D). This finding suggests that taste identity is encoded mainly by power and phase in the delta-frequency range (1-4 Hz), and therefore, the subsequent analysis focused solely on this frequency.

**Figure 1.**
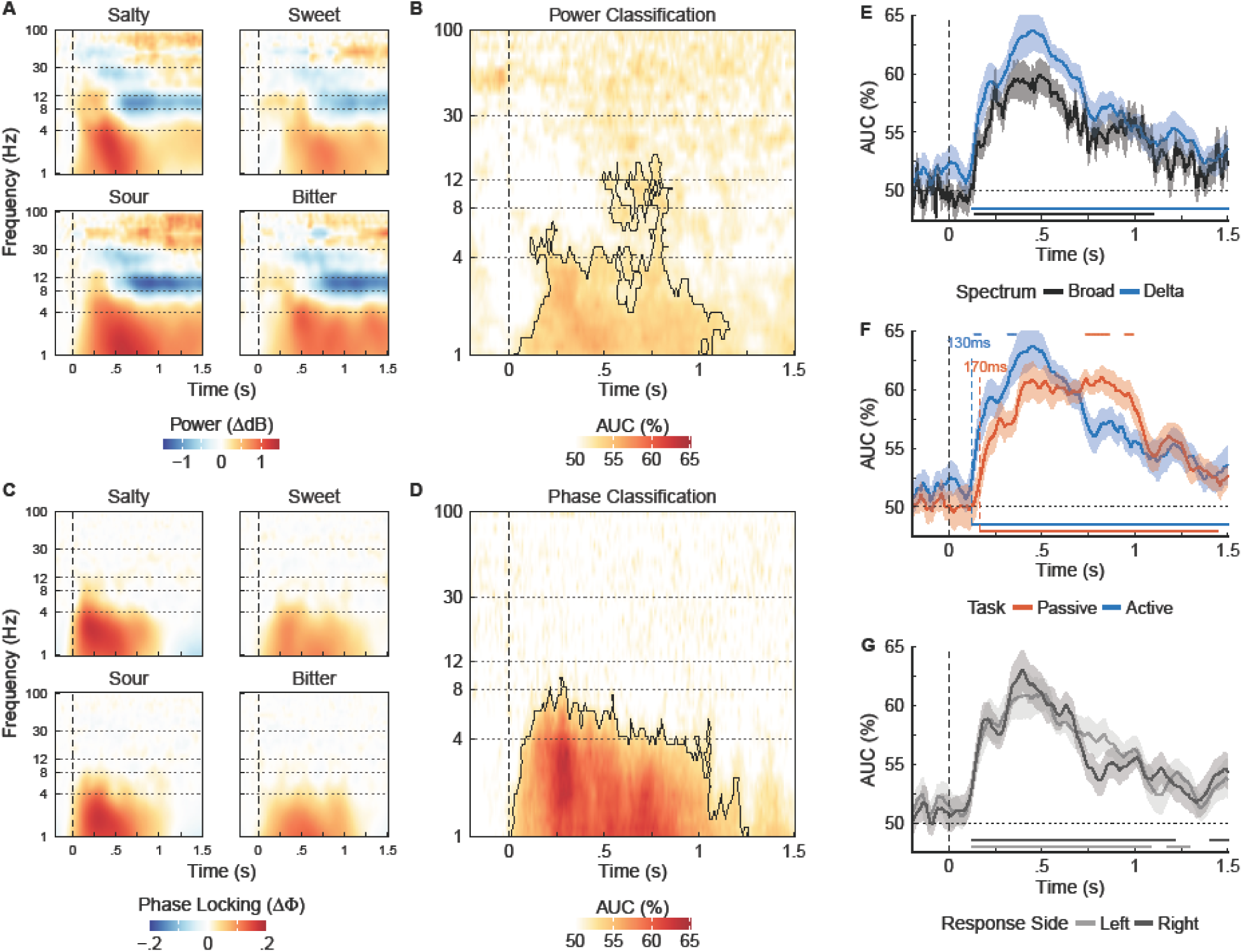
Taste information is encoded by delta oscillations. A) Spectral power estimates obtained via continuous wavelet transformation of the EEG recordings for frequencies from 1 to 100 Hz and averaged across 64 electrodes and 16 participants. The power represents the amplitude of the signal. Increases (warm colours) and decreases (cold colours) are relative to the baseline (for visualization). Vertical dashed line: stimulus onset; horizontal dotted lines: transitions between established frequency bands (1-4 Hz: delta, 4-8 Hz: theta, 8-12 Hz: alpha, 12-30 Hz: beta, 30-100 Hz: gamma; cf. Herrmann et al., 2005). B) SVM classifiers were trained at each time point and frequency step to decode the four tastes given the (non-normalized) spectral power in the 64-electrode space. Performance is displayed as the AUC (50% corresponds to the chance level) averaged across participants (black contour lines indicate cluster-corrected significance). C) Inter-trial coherence (ITC) is calculated from phase estimates obtained via continuous wavelet transformation for frequencies from 1 to 100 Hz. The ITC expresses the extent of phase synchronization across trials (remainder as in A). D) SVM classifiers were trained at each time point and frequency step to decode the four tastes given the phases in radians in the 64-electrode space (remainder as in B). E) Comparison of taste response-pattern decoding between the broad-band and delta signal (1-4 Hz). SVM classifiers were trained at each time point to decode the four tastes, given the unfiltered (black) or FIR filtered (blue) electrophysiological recordings from 64 electrodes (solid lines: mean AUC across trials and participants; surrounding shaded regions ±1 SEM; colour-coded horizontal lines above the x-axis indicate above-chance performance; horizontal dotted line: theoretical chance level of 50%). F) Comparison between the speeded, active task of the current study and a delayed, passive task (Crouzet et al., 2015). Colour-coded stars at the top indicate significantly different classification performance between active and passive task; colour-coded dashed vertical lines indicate task-specific starting times of significant multi-taste discrimination (remainder as in E). G) Motor-control task. Response mappings were separated for the training and test set, i.e. the decoder was trained on trials with a left button press, and tested on trials with a right button press (and vice versa). Successful classification demonstrates that motor-related activity is irrelevant to the multi-taste discrimination (remainder as in E).

To optimize temporal precision, we decoded taste identity information from the electrical recordings after band-pass zero-phase FIR filtering (0.5-4.5Hz±1; order: 330) to isolate delta activity from other frequencies. We identified the earliest point of significant above-chance classification performance at 130 ms (AUC = 53.0%±1.4, *p* = .005); it lasted until the end of the epoch (AUC_max_ = 63.7%±1.7, *p* < .0001, at 450 ms; Fig. 1E). Moreover, in contrast to the unfiltered recordings, the delta-encoded signal yielded higher classification performance throughout the epoch, suggesting an overall improved signal-to-noise ratio (AUC_maxdiff_ = 5.3% at 460 ms, *p* = .007; cf. Fig. 1E). Notably, taste patterns appeared to emerge earlier during this active, speeded taste-detection task than in a previously reported study, in which participants passively tasted and gave delayed taste categorization responses (Crouzet et al., 2015). To test this, we applied the multi-class decoding analysis reported here using the FIR filtered electrophysiological recordings obtained during the passive tasting task and compared the findings (Fig. 1F). We found that classification performance reached significance 40 ms later in the passive task, compared to the active one (at 170 ms, AUC = 52.6%±1.4, *p* = .022). The classification performance of the active task was significantly higher than the passive task during the periods from 140 to 170 ms and from 320 to 360 ms (all *p* < .0001; AUC_maxdiff_ = 4.7% at 160 ms). In contrast, the classification performance of the passive task was significantly higher in later periods from 740 to 860 ms and from 950 to 990 ms (all *p* < .0001; AUC_maxdiff_ = 4.7% at 790 ms). Together, these findings indicate that the taste response pattern is susceptible to task demands.

To exclude the possibility that the classifiers were using motor-related activation to solve the multi-taste discrimination, we reiterated the procedure by carefully separating the response sides, so that classifiers were trained on patterns from trials with right-sided responses and tested on patterns of trials with left-sided responses, and vice versa. We found highly similar performance curves for classifiers trained on right- and left-sided responses with above-chance classification starting at 130 ms for most of the time range for both response sides (training left, testing right: AUC_onset_ = 53.6%±1.0, *p* = .022, AUC_max_ = 61.0%±1.3 at 520 ms; training right, testing left: AUC_onset_ = 53.6%±1.1, *p* = .026; AUC_max_ = 63.0%±1.7 at 400 ms; Figure 1G). Importantly, the performance curves for response-side and taste decoding were highly similar as well (compare Figures 1E and 1G). The interchangeability of the response side topographies during classification confirms that the taste discriminability within the delta spectrum was independent of motor activation.

### The emergence of taste-evoked delta activity is predictive of taste detection responses

The above findings indicate delta activation as an electrophysiological signature of taste-information coding. To test whether there was a systematic link between these patterns and taste detection times, we computed multi-level linear mixed models between the onset of delta-encoded taste information and response times. We used SVM classifiers to discriminate each trial from its baseline period water rinse, which closely matched the taste-detection task that participants performed in a given trial (Fig. 2A). We found a highly significant positive relationship between the onsets of taste-pattern decoding and taste detection response times (*β* = .32±.03, *χ*^2^_(1,*N*=2990)_ = 16.87, *p* < .0001; Fig. 2B) for each of the four tastes: salty (*M*_on_ = 185 ms, *M*_RT_ = 643 ms, = *β*=.35), sour (*M*_on_ = 196 ms, *M*_RT_ = 680 ms, *β* = .34), sweet (*M*_on_ = 242 ms, *M*_RT_ = 845 ms, *β* = .30), and bitter (*M*_on_ = 244 ms, *M*_RT_ = 798 ms, *β* = .31). This finding indicates a functional relevance of this activation pattern for taste detection behavior, such that faster taste-pattern emergence can be linked to faster taste-detection times.

**Figure 2.**
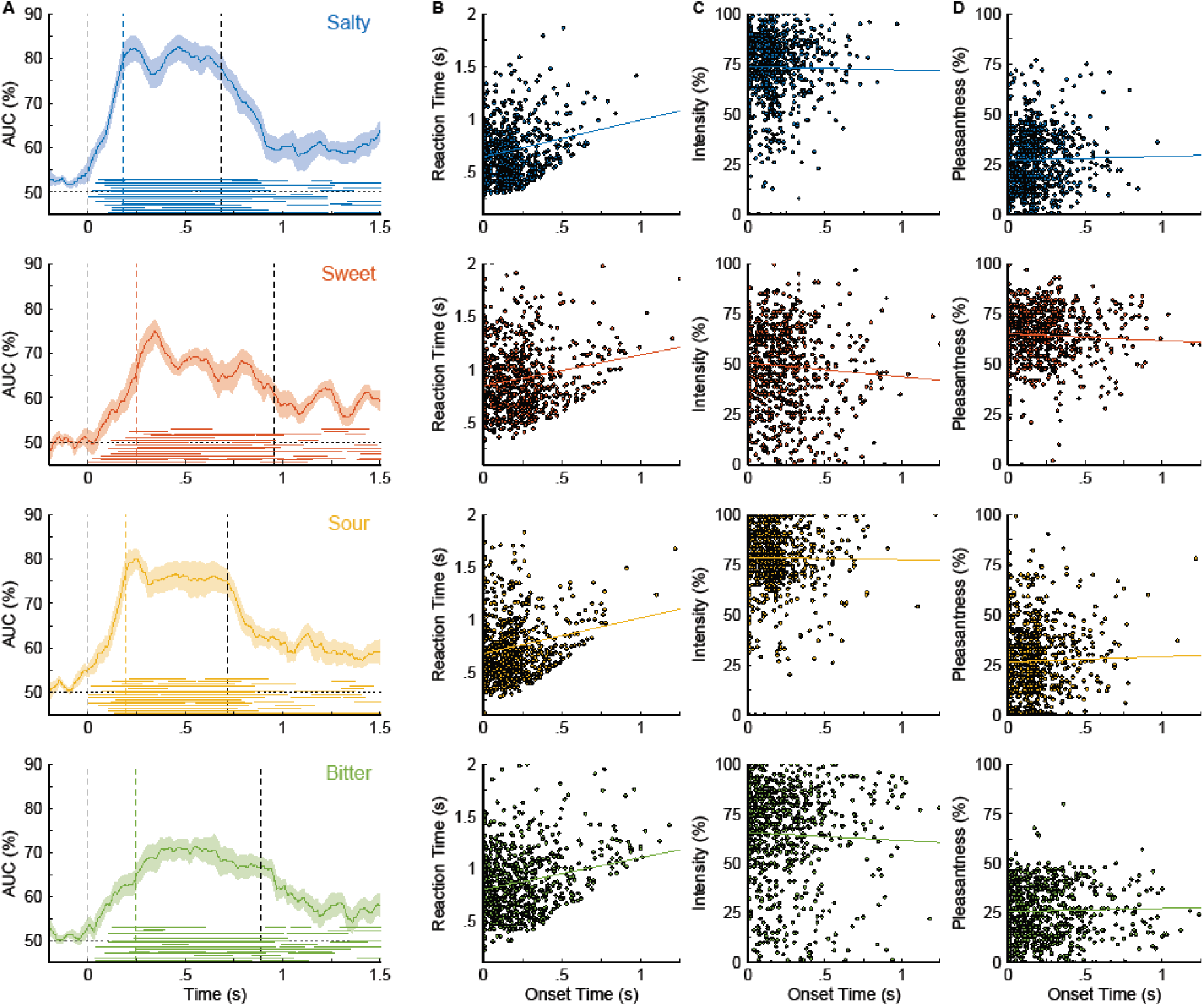
The emergence of the delta-encoded electrophysiological taste signature is predictive of taste responses. A) SVM classifiers were trained within participants at each time point on the patterns of N-1 trials, in order to decode taste and water in the left-out trial (each trial was matched with its water baseline average). This classification task closely corresponds to the taste-detection task participants performed (press as soon as you taste something). Colour-coded solid lines: mean AUC across participants, surrounding shaded regions ± 1 SEM; colour-coded horizontal lines above the x-axis indicate periods of above-chance performance per participant; colour-coded dashed vertical lines: average onset time; black dashed vertical lines: average response time; grey vertical dashed line: stimulus onset; black horizontal dotted line: theoretical chance level of 50%. B) Correlation plots between the onsets of pattern decoding (determined as the earliest point at which a trial has been classified successfully for at least 100 ms) and taste-detection response times, taste-intensity rating scores (C), and taste-pleasantness rating scores (D). Colour-coded solid lines indicate the taste-specific influence of decoding onset from the multi-level linear mixed regression fits.

To determine whether the observed delta activity was encoding taste identity rather than taste intensity or valence, we computed analogous mixed linear models using the intensity and pleasantness ratings as dependent variables. We observed no significant relationship between the onsets of taste-pattern decoding (quantified in ms) and intensity ratings (*β* = –.004±.002, *χ*^2^_(1,*N*=2984)_ = 2.33, *p* = .127; *M*_salty_ = 74, *M*_sour_ = 79, *M*_sweet_ = 51, *M*_bitter_ = 66; Fig. 2C), and pleasantness ratings (*β* = .001±.002, *χ*^2^_(1,*N*=2985)_ = 0.15, *p* = .699; *M*_salty_ = 27, *M*_sour_ = 27, *M*_sweet_ = 65, *M*_bitter_ = 26; Fig. 2D). Together, these results suggest that delta activity encodes information of taste identity rather than other taste features.

## DISCUSSION

The fundamental role of coherent rhythmic activity in neuronal communication has been well established for perception and cognition across sensory systems and species (von Stein et al., 2000; Engel et al., 2001; Varela et al., 2001; Fries, 2005; Buzaki, 2006). Here, we identified for the first time delta activity (1-4 Hz) as a distinct electrophysiological signature of gustatory processing in the human brain. This finding adds to recent evidence that highlights the relevance of slow cortical rhythms in the coding of taste-specific information in the rodent gustatory cortex (<1.5 and 4-5 Hz; Pavao et al., 2014) and odor-specific content in the human olfactory cortex (4-8 Hz; Jiang et al., 2017).

Primate taste processing involves neural computations in a widespread network, including the insula (primary gustatory area) and the orbitofrontal cortex (secondary gustatory area) (Pritchard and Di Lorenzo, 2015), which further link to subcortical structures regulating reward and feeding behavior (Katz and Sadacca, 2011), as well as somatosensory and visceral areas (Katz et al., 2002). Given that the length of an oscillatory cycle determines the range and timescale of cortical integration, such that slower oscillations bridge longer distances and accomplish more complex computations (Engel et al., 2010; Harmony, 2013), our findings of delta-encoded taste information may indicate the mechanism by which the human brain communicates within such a spatially distributed and highly interconnected system. Delta oscillations have been found to originate in the primate insula (Mesulam and Mufson, 1982), and they play a role in food-related arousal such as craving and reward (cf. Knyazev, 2007, 2012). We supplement this knowledge with evidence of a novel role of delta activity in coding taste information in the human brain. Here, the slow-wave activity subsides with the offset of stimulation. Given the successful and improved classification of the taste quality information from the spectral signatures after attenuation of non-delta frequencies, we interpret this finding in accordance with the literature on slow rhythmic activity, which has been suggested to subserve the sequencing of a sensory experience into discrete processing entities (Colgin, 2013; Wilson et al., 2015). Accordingly, each processing cycle coordinates through its phase the activation of cell populations that represent specific experiential aspects, whereas the cycle sequence links the individual segments to maintain the integrity of the experience. Through phase coherence (phase-coordinated activity), phase-segregated information bits become accessible to distant brain areas. Our findings suggest that delta activity contributes the candidate frequency with which such distinct information packages are transmitted from the same taste episode across the long spatial trajectory of the taste network. This interpretation will benefit from additional electrocorticographical data and modelling. In line with the notion that phase coherence coordinates neuronal activation, we found that the phase encoded more information than its amplitude. More than a means of coordinating clearly defined bits of information, in sensory cortices, the functional role of slow oscillations has been proposed as an internal reference frame which stabilizes the encoding of natural stimuli, thereby fostering perceptual robustness (Panzeri et al., 2014). This notion is especially intriguing for the chemical senses, for which sensory representations have to be formed in environments with a high degree of (temporal) uncertainty (cf. Jiang et al., 2017). Among other factors, the gustatory system encounters noise as sensory inputs originate from different spatial locations within the oral cavity over an extended period of mastication and swallowing with no precisely timed stimulus. Indeed, the action potentials of (chemo-)sensory neurons are much more precisely timed than the chemical stimulation they respond to. When these action potentials are summed over long time windows, sensory information is at risk to be degraded or lost. Accordingly, an oscillatory reference frame may act as a pacemaker, which helps decode the incoming information from finely timed spike patterns (Panzeri et al., 2014). Thus, the gustatory system, like the olfactory, would benefit particularly from slow rhythmic activity as a means of attaining perceptual robustness.

To determine whether delta-encoded taste information is in fact used for perceptual decisions, we correlated the onset of delta activity with the behavioral response time in each trial. This onset predicted when participants detected a given taste, such that a more rapid pattern-emergence corresponded to faster response behavior. Importantly, we excluded the possibility of decoding performance being confounded by motor signals through careful cross-response side validation. This apparent neural-behavioral link aligns well with the aforementioned notion that slow oscillations are not just a by-product of network processing, but have functional significance for the perceptual integration and temporal alignment of sensory inputs through coherence (e.g. via delay-based synchrony detection). Moreover – although the units by which neurons encode information are precisely timed spike patterns within a range of a few milliseconds – the behaviorally relevant information may very well extend over hundreds of milliseconds. Therefore, slow rhythmic patterns can encode information that is not available in local spike counts (Kayser et al., 2012), and thus enable other brain structures to extract relevant sensory information which informs decisions or concomitant perceptual states (Wilson et al., 2015). Indeed, the present data provide evidence of such a link between the timing of delta activity and subsequent behavior.

To test whether the timing of the earliest sensory taste-response is independent of behavioral demands, we compared the decoding performance between two different tasks: a fast paced, speeded taste-detection task (reported here) and a slow paced, delayed taste-categorization task (Crouzet et al., 2015). We found earlier and stronger delta-encoded taste discriminability in the speeded and more sustained taste discriminability in the delayed task, suggesting that neural taste responses are contingent upon behavioral goals. Task-dependent latency differences in event-related potentials have been observed previously for other modalities, although it remains unclear whether these effects reflect actual shifts in the availability of information. The observed flexibility in taste-information coding accords well with the observation that gustatory cortical activity varies greatly with expectation and focus of attention in rodents (Fontanini and Katz, 2009). Our findings show that the degree and speed at which the taste system categorizes stimuli is likely to be tuned to situational demands, thereby enabling adaptive behaviour.

In summary, we present evidence of activation in the delta-frequency range as an electrophysiological signature of taste processing in humans. We show that taste-specific content can be discerned from these signatures, which predict perceptual decisions and align flexibly with task requirements.

## Conflict of Interest

The authors declare no competing financial interests.

## Acknowledgements

This work was supported by NutriAct–Competence Cluster Nutrition Research Berlin-Potsdam funded by the Federal Ministry of Education and Research (FKZ: 01EA1408A-G) granted to KO.

## References

Accolla R, Bathellier B, Petersen CC, Carleton A (2007) Differential spatial representation of taste modalities in the rat gustatory cortex. J Neurosci 27:1396–1404.

Bujas Z, Szabo S, Ajdukovic D, Mayer D (1989) Individual gustatory reaction times to various groups of chemicals that provoke basic taste qualities. Percept Psychophys 45:385–390.

Busch NA, Dubois J, VanRullen R (2009) The phase of ongoing EEG oscillations predicts visual perception. The Journal of neuroscience : the official journal of the Society for Neuroscience 29:7869–7876.

Buzaki G (2006) Rhythms of The Brain: Oxford University Press.

Carleton A, Accolla R, Simon SA (2010) Coding in the mammalian gustatory system. Trends Neurosci 33:326–334.

Chandrashekar J, Hoon MA, Ryba NJP, Zuker CS (2006) The receptors and cells for mammalian taste. Nature 444:288–294.

Chen X, Gabitto M, Peng Y, Ryba NJP, Zuker CS (2011) A gustotopic map of taste qualities in the mammalian brain. Science (New York, NY) 333:1262–1266.

Colgin LL (2013) Mechanisms and functions of theta rhythms. Annu Rev Neurosci 36:295–312.

Crouzet SM, Busch NA, Ohla K (2015) Taste quality decoding parallels taste sensations. Current Biology 25(7):890–896.

Delorme A, Makeig S (2004) EEGLAB: an open source toolbox for analysis of single-trial EEG dynamics including independent component analysis. Journal of Neuroscience Methods 134:9–21.

Engel AK, Fries P, Singer W (2001) Dynamic predictions: oscillations and synchrony in top-down processing. Nat Rev Neurosci 2:704–716.

Engel AK, Friston K, Kelso JAS, König P, Kovács I, MacDonald III A, Miller EK, Phillips WA, Silverstein SM, Tallon-Baudry C, Triesch J, Uhlhaas P (2010) Coordination in Behavior and Cognition. In: Dynamic Coordination in the Brain: From Neurons to Mind (von der Malsburg C, Phillips WA, Singer W, eds), pp 267–300: MIT Press.

Fan R, Chang K, Hsieh C (2008) LIBLINEAR: A library for large linear classification. The Journal of Machine Learning Research 9:1871–1874.

Fontanini A, Katz DB (2009) Behavioral modulation of gustatory cortical activity. Ann N Y Acad Sci 1170:403–406.

Fries P (2005) A mechanism for cognitive dynamics: neuronal communication through neuronal coherence. Trends in cognitive sciences 9:474–480.

Fries P (2015) Rhythms for Cognition: Communication through Coherence. Neuron 88:220–235.

Grootswagers T, Wardle SG, Carlson TA (2017) Decoding Dynamic Brain Patterns from Evoked Responses: A Tutorial on Multivariate Pattern Analysis Applied to Time Series Neuroimaging Data. Journal of cognitive neuroscience 29:677–697.

Halpern BP (1986) Constraints imposed on taste physiology by human taste reaction time data. Neurosci Biobehav Rev 10:135–151.

Hand DJ, Till RJ (2001) A simple generalisation of the area under the ROC curve for multiple class classification problems. Machine learning 45:171–186.

Harmony T (2013) The functional significance of delta oscillations in cognitive processing. Frontiers in integrative neuroscience 7:83.

Herrmann CS, Grigutsch M, Busch NA (2005) EEG oscillations and wavelet analysis. In: Event-related Potentials: A Methods Handbook (Handy TC, ed), pp 229–259. Cambridge, MA: MIT Press.

Jas M, Engemann DA, Bekhti Y, Raimondo F, Gramfort A (2017) Autoreject: Automated artifact rejection for MEG and EEG data. In, 12 Jul 2017 Edition, pp 1–24. arXiv.

Jiang H, Schuele S, Rosenow J, Zelano C, Parvizi J, Tao JX, Wu S, Gottfried JA (2017) Theta Oscillations Rapidly Convey Odor-Specific Content in Human Piriform Cortex. Neuron 94:207–219 e204.

Jones LM, Fontanini A, Sadacca BF, Miller P, Katz DB (2007) Natural stimuli evoke dynamic sequences of states in sensory cortical ensembles. Proceedings of the National Academy of Sciences of the United States of America 104:18772–18777.

Kahana MJ (2006) The cognitive correlates of human brain oscillations. The Journal of neuroscience : the official journal of the Society for Neuroscience 26:1669–1672.

Katz DB, Sadacca BF (2011) Taste. In: Neurobiology of Sensation and Reward (Gottfried JA, ed). Boca Raton, FL, USA: CRC Press/Taylor & Francis.

Katz DB, Nicolelis MA, Simon SA (2002) Gustatory processing is dynamic and distributed. Current opinion in neurobiology 12:448–454.

Kayser C, Ince RA, Panzeri S (2012) Analysis of slow (theta) oscillations as a potential temporal reference frame for information coding in sensory cortices. PLoS computational biology 8:e1002717.

Klimesch W (2012) alpha-band oscillations, attention, and controlled access to stored information. Trends in cognitive sciences 16:606–617.

Knyazev GG (2007) Motivation, emotion, and their inhibitory control mirrored in brain oscillations. Neuroscience and biobehavioral reviews 31:377–395.

Knyazev GG (2012) EEG delta oscillations as a correlate of basic homeostatic and motivational processes. Neuroscience and biobehavioral reviews 36:677–695.

Koepsell K, Wang X, Hirsch JA, Sommer FT (2010) Exploring the function of neural oscillations in early sensory systems. Frontiers in neuroscience 4:53.

Kriegeskorte N (2011) Pattern-information analysis: from stimulus decoding to computational-model testing. NeuroImage 56:411–421.

Lopes da Silva F (1991) Neural mechanisms underlying brain waves: from neural membranes to networks. Electroencephalography and clinical neurophysiology 79:81–93.

Makeig S, Jung TP, Bell AJ, Ghahremani D, Sejnowski TJ (1997) Blind separation of auditory event-related brain responses into independent components. Proc Natl Acad Sci U S A 94:10979–10984.

Maris E, Oostenveld R (2007) Nonparametric statistical testing of EEG- and MEG-data. Journal of neuroscience methods 164:177–190.

Mesulam MM, Mufson EJ (1982) Insula of the old world monkey. III: Efferent cortical output and comments on function. The Journal of comparative neurology 212:38–52.

Mognon A, Jovicich J, Bruzzone L, Buiatti M (2011) ADJUST: An automatic EEG artifact detector based on the joint use of spatial and temporal features. Psychophysiology 48:229–240.

Panzeri S, Ince RA, Diamond ME, Kayser C (2014) Reading spike timing without a clock: intrinsic decoding of spike trains. Philosophical transactions of the Royal Society of London Series B, Biological sciences 369:20120467.

Pavao R, Piette CE, Lopes-dos-Santos V, Katz DB, Tort ABL (2014) Local field potentials in the gustatory cortex carry taste information. The Journal of neuroscience : the official journal of the Society for Neuroscience 34:8778–8787.

Platt J (1999) Probabilistic outputs for support vector machines and comparisons to regularized likelihood methods. Advances in large margin classifiers 10:61–74.

Pritchard TC, Di Lorenzo PM (2015) Central Taste Anatomy and Physiology of Rodents and Primates. In: Handbook of Olfaction and Gustation (Doty RL, ed), pp 701–726. Hoboken, NJ, USA: John Wiley & Sons, Inc.

R-Core-Team (2017) R: A Language and Environment for Statistical Computing. In. Vienna, Austria: R Foundation for Statistical Computing.

Rosenstein D, Oster H (1988) Differential facial responses to four basic tastes in newborns. Child Dev 59:1555–1568.

Salenius S, Hari R (2003) Synchronous cortical oscillatory activity during motor action. Current opinion in neurobiology 13:678–684.

Stapleton JR, Lavine ML, Wolpert RL, Nicolelis MA, Simon SA (2006) Rapid taste responses in the gustatory cortex during licking. The Journal of neuroscience : the official journal of the Society for Neuroscience 26:4126–4138.

Tallon-Baudry C (2003) Oscillatory synchrony and human visual cognition. Journal of physiology, Paris 97:355–363.

Varela F, Lachaux JP, Rodriguez E, Martinerie J (2001) The brainweb: phase synchronization and large-scale integration. Nature reviews Neuroscience 2:229–239.

von Stein A, Chiang C, Konig P (2000) Top-down processing mediated by interareal synchronization. Proceedings of the National Academy of Sciences of the United States of America 97:14748–14753.

Ward LM (2003) Synchronous neural oscillations and cognitive processes. Trends in cognitive sciences 7:553–559.

Wilson MA, Varela C, Remondes M (2015) Phase organization of network computations. Current opinion in neurobiology 31:250–253.

